# Why are SNAREpins rod-shaped?

**DOI:** 10.1101/2021.06.23.449668

**Authors:** Jin Zeng, Zachary McDargh, Dong An, Ben O’Shaughnessy

## Abstract

SNARE proteins are the core components of the cellular machineries that fuse membranes for neurotransmitter or hormone release and other fundamental processes. Fusion is accomplished when SNARE proteins hosted by apposing membranes form SNARE complexes called SNAREpins, but the mechanism of fusion remains unclear. Computational simulations of SNARE-mediated membrane fusion are challenging due to the millisecond timescales of physiological membrane fusion. Here we used ultra-coarse-grained (UCG) simulations to investigate the minimal requirements for a molecular intracellular fusogen, and to elucidate the mechanisms of SNARE-mediated fusion. We find fusion by simple body forces that push vesicles together is highly inefficient. Inter-vesicle fusogens with different aspect ratios can fuse vesicles only if they are rodlike, of sufficient length to clear the fusogens from the fusion site by entropic forces. Simulations with rod-shaped SNAREpin-like fusogens fused 50-nm vesicles on ms timescales, driven by entropic forces along a reproducible fusion pathway. SNARE-SNARE and SNARE-membrane entropic forces cleared the fusion site and pressed the vesicles into an extended contact zone (ECZ), drove stalk nucleation at the high curvature ECZ boundary, and expanded the stalk into a long-lived hemifusion diaphragm in which a simple pore completed fusion. Our results provide strong support for the entropic hypothesis of SNARE-mediated membrane fusion, and implicate the rodlike structure of the SNAREpin complex as a necessity for entropic force production and fusion.

## Introduction

Fundamental processes such as neurotransmitter (NT) or hormone release rely on regulated exocytosis, when a signal, usually Ca^2+^, triggers fusion of docked secretory vesicles to the plasma membrane (PM), releasing signaling or other bioactive molecules via the fusion pore. Evoked exocytotic release is accomplished by specialized cellular machineries. SNARE proteins are the core of these fusion machineries. Vesicle-associated v-SNAREs form SNAREpin complexes with PM-associated t-SNAREs, pulling the membranes together for fusion (1, 2).

How the ~ 10 nm long rod-shaped SNARE complexes (“SNAREpins”) accomplish fusion is not established. SNARE-mediated fusion is often thought driven by some fraction of the ~60 *k*_*B*_*T* zippering energy of SNARE complex formation(3, 4), but how this energy is transduced into the membranes is unexplained. Experiments suggest fusion requires a certain minimum number of SNAREpins, but the reported requirements vary widely. In vivo experimental evidence suggests the number of SNAREpins required for fusion ranges from 2-8 (5–8). In vitro, 1-3 SNAREpins were sufficient to fuse small unilamellar vesicles (SUVs) (9, 10), but 5-10 SNAREpins were required to fuse SUVs with supported bilayers (11, 12). Interestingly, large unilamellar vesicle (LUV) fusion required up to ~ 30 SNAREpins (13).

SNAREpins could use zippering fores for fusion, by the linker domains (LDs) connecting the Syntaxin and VAMP SNARE motifs to TMDs (14, 15) exerting bending torques on the membrane (16). However, evidence from NMR and EPR suggests the LDs are unstructured and flexible (17–21). In functional studies with short flexible sequences inserted into the LDs fusion is progressively impaired rather than completely abolished (22–24), though a recent study found that fusion could be abolished depends on the precise insertion location (25).

Many computational molecular dynamics (MD) studies have sought to elucidate the mechanisms, but SNARE-mediated fusion was rarely observed. Fusion timescales appear inaccessible to all-atom (AA) approaches (16, 26). However SNARE-driven fusion was achieved using the coarse-grained (CG) MARTINI force field approach (27, 28). In one study planar membranes and nanodiscs fused via a minimal hemifused connection, the stalk, and then an inverted micelle intermediate (27). In another groundbreaking study, 20 nm pure PE lipid vesicles fused via stalk, inverted micelle and enlarged hemifusion diaphragm (HD) intermediates (28). Both studies achieved fusion in < 1 μs, within reach of MARTINI, a surprisingly short timescale given that electrophysiological measurements of synaptic vesicle release rates suggest ~ ms fusion times (29–32). However, on one case fusion was favored by small vesicle sizes and unphysical lipid composition (28), and fusion in both studies relied on stiff LDs (27, 28).

We previously used simulations with highly coarse-grained (CG) SNAREs and continuous membrane surfaces to access physiological ms timescales (33, 34). These studies suggest that SNAREs fuse membranes by entropic forces, a radically different mechanism to the hypothesized mechanisms based on zippering energy or LD stiffness. In simulations with SNARE proteins bridging a vesicle and planar membrane, the waiting time for the membrane energy to attain the fusion energy threshold was measured Any number of SNAREs catalyzed fusion, but more SNAREs gave faster fusion (33, 34). Entropic forces organized the fully assembled SNAREpins into a ring and pulled the membranes together for fusion.

TMD-mediated membrane thinning may promote fusion (35, 36). Experiments and CG simulations showed that TMDs induce can locally thin membranes due to mismatch in the length of the hydrophobic regions of the TMDs and the membrane, proposed to segregate SNARE proteins into microdomains in cells. Another MARTINI study found TMD-induced membrane thinning lowers the activation energy for stalk formation (36).

Considerable experimental evidence suggests extended contact and hemifused states lie on the pathway to fusion. In protein-free calcium-driven studies, GUVs adhered and then formed an HD connection which subsequently ruptured for fusion (37), a pathway that was quantiatively reproduced by a mathematical model (37–39). Extended contact and hemifused states have been observed in SNARE-reconstituted vesicle-vesicle fusion systems (40, 41). HDs have also been observed in live cells. Dense-core vesicles made hemifused connections featuring and HD with the plasma membrane of chromaffin and pancreatic β cells (42).

Here we simulated fusion of ultra-coarse-grained (UCG) explicit lipid membranes by UCG fusogens of different classes, building up to SNARE-like fusogens. We sought to establish the minimal requirements for a molecular intracellular fusogen, and to elucidate the mechanisms of SNARE-mediated fusion. We first fused membranes by simple body forces that push vesicles together (43), which we find is highly inefficient. Inter-vesicle fusogens with different aspect ratios were then simulated, showing that fusion requires rodlike shapes of sufficient length to clear the fusogens from the fusion site by entropic forces. Finally, simulations with rod-shaped SNARE-like fusogens were able to fuse 50-nm vesicles on ms timescales, driven by entropic forces. The fusion pathway was reproducible: SNARE-SNARE and SNARE-membrane entropic forces cleared the fusion site and pressed the vesicles into an extended contact zone (ECZ), drove a stalk to nucleate at the high curvature ECZ boundary, and drove stalk expansion into a long-lived hemifusion diaphragm (HD) in which a simple pore completed fusion. These results are consistent with our earlier simulations using simple continuous membranes (33, 34), but now articulate the deformed state of the adhered vesicles resulting from the entropic forces that press them together. Our results provide strong support for the entropic hypothesis of SNARE-mediated membrane fusion.

## Results

### Model

To determine the essential features of productive molecular fusogens, we performed molecular dynamics (MD) simulations and studied whether either externally applied forces or model molecular fusogens could fuse two vesicle membranes. Since the fusion timescale in neurotransmitter release is sub-millisecond (44, 45), far beyond what is achievable with atomistic or widely used MARTINI models, we used coarse-grained (CG) representations of phospholipids and molecular fusogens to test for possible fusion events on these timescales.

#### Ultra-coarse-grained representation of phospholipids

We employed the coarse-grained Cooke-Deserno lipid model in our simulations (46–48). In this model, the phospholipid is represented by linearly connected beads consisting of one head bead (h) and three tail beads (t) (Fig.1A), which interact through an implicit-solvent force field. For interaction details, please see SI. The CG length unit σ is set to 0.88nm (46–48) by equating the CG lipid bilayer thickness (~ 5.7σ) with the experimentally observed membrane thickness of 5 nm for pure phosphatidylcholine (49). The CG energy unit ϵ is set to 0.6*k*_*B*_*T* (46–48), and the attractive potential decay range *w*_*c*_ is set to 1.6σ (46–48).

All lipid beads are defined to have mass *m* = 1 (46–48). The simulation time step Δ*t* is determined as 0.068 ns (46–48) by equating the measured lateral lipid diffusivity 8.8 × 10^−5^ *σ*^2^/Δ*t* from simulations with the experimental measurement of 1μm^2^/s in a mixed lipid membrane (DOPC/SM/CHOL) (50).

#### SNARE-like fusogen model

The synaptic SNARE complex is a coiled-coil of four *α*-helices (51, 52), where two helices are from SNAP-25, and the other two helices come from VAMP and Syntaxin (Stx), respectively. VAMP and Stx are anchored to membranes by their transmembrane domains (TMDs), which are connected to the SNARE motif by linker domains (LDs) (Fig. 1C) (52). We constructed a minimal model fusogen designed to mimic the most basic features of the SNARE complex; the structure of these SNARE-like fusogens was based on the neuronal SNARE complex crystal structure (52), shown in Fig.1 D, E. We represent the coiled-coil SNARE motif as a rigid body with a length *L*_SNARE_ = 10 nm and a width *D*_SNARE_ = 2 nm consisting of 9 linearly placed beads. The *α* -helical TMDs of VAMP and Stx are each represented by a rigid body with a length *L*_TMD_ = 3 nm and a width *D*_TMD_ = 1 nm, consisting of 3 linearly placed beads (Fig.1 D).

**Figure 1.**
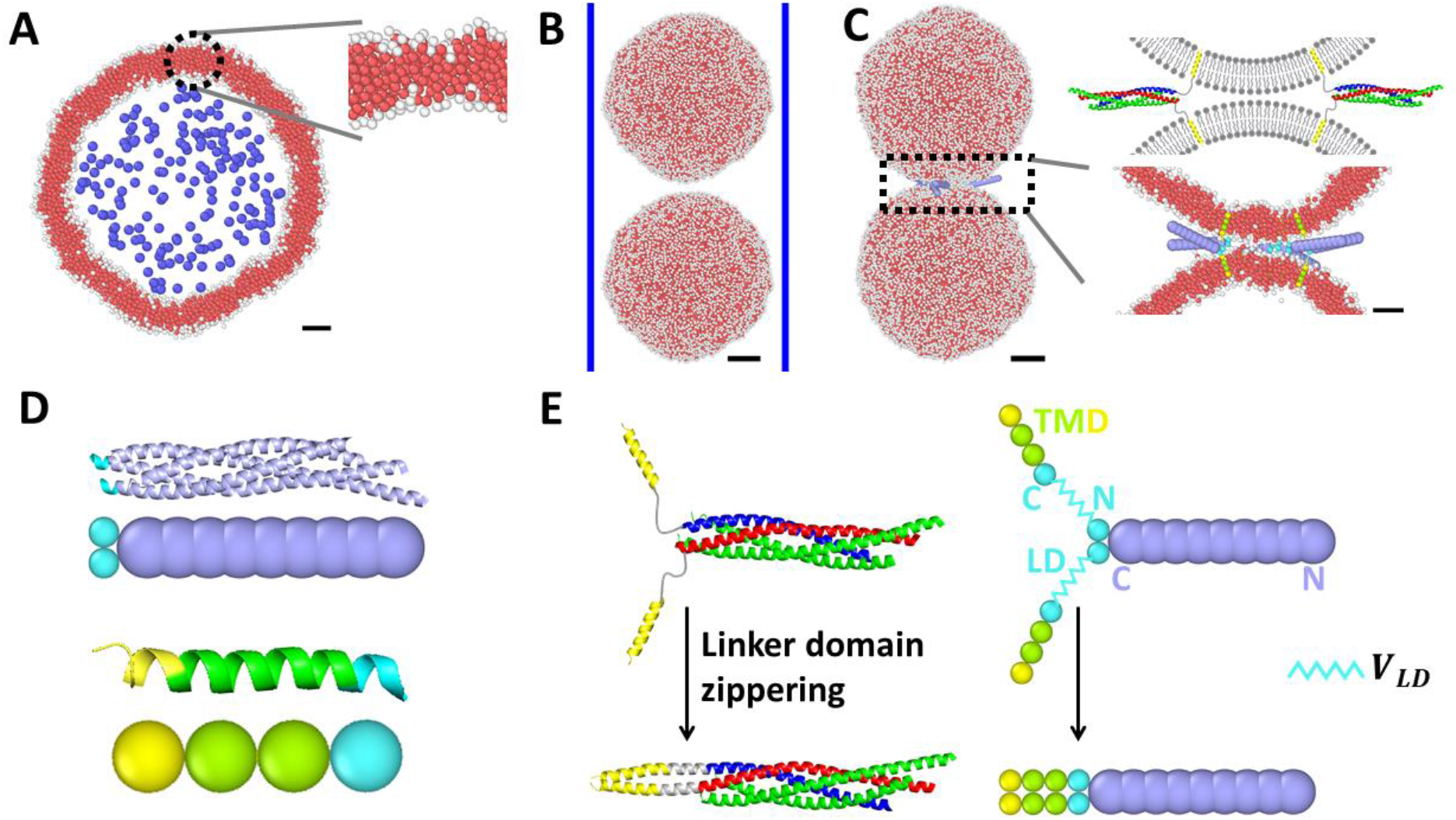
Coarse-grained model of phospholipid vesicles and SNARE complexes. (A) Cross-section of a simulated ~50 nm diameter CG phospholipid vesicle, in which lipid head beads are shown white and lipid tail beads red. Solvent beads (blue) in the vesicle lumen are used to maintain the membrane tension. Scale bars: 10 nm. (B) Side view of the simulation initial condition in the absence of SNARE-like fusogens, in which two ~50 nm diameter vesicles are placed within a cylinder (blue lines) and pressed together by forces applied to every bead. Scale bars: 10 nm. (C) Simulation of SNARE-mediated fusion. Left: Side view of the simulation initial condition, in which 6 SNARE-like fusogens connect two ~50 nm diameter vesicles, with lipids overlapping the TMDs deleted. Scale bars: 10 nm. Top right: Schematic of two trans-SNARE complexes connecting two vesicle membranes. VAMP is colored blue; Stx is colored red, and SNAP-25 is colored green. VAMP and Stx are anchored to a vesicle membrane by *α*-helical TMDs (yellow)(PDB ID: 3HD7) (52). Bottom right: zoomed in view of the 6 SNARE-like fusogens connecting two vesicles. Scale bars: 5 nm. (D) Top: SNARE complex and the coarse-grained model. Bottom: TMD and the coarse-grained model. The TMD consists of two hydrophobic beads (green) and one hydrophilic bead (yellow). An LD C-terminus bead (cyan) attaches to the TMD rigid body. (E) Top: the neuronal SNARE complex and the CG SNARE complex with unzippered LDs. Bottom: zipppered SNARE complex.

Experiments using optical tweezers to unzipper the neuronal SNARE complex found that SNARE complex unzippering requires a ~18 pN force, and releases ~65 *k*_*B*_*T* in zippering free energy (3). The SNARE zippering free energy landscape is approximately linear near the SNARE motif C-terminus (3), i.e. the zippering force is constant. Thus we used a linear zippering potential, tapering harmonically to zero,

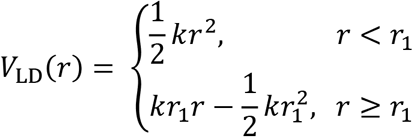

with *r*_1_ = 0.1nm and *k* = *T*_zip_/*r*_1_, where *T*_zip_ = 18pN is the zippering force. To incorporate the LD in the model, one bead with a diameter *D*_TMD_ located at the LD C-terminus interacts with a point at the LD N-terminus via the potential *V*_LD_(*r*) where *r* is the LD length (Fig. 1E).

Non-bonded interactions involving SNARE-like fusogen beads were set similar to the Cooke-Deserno lipid model. We used the WCA potential to define each bead size. In addition to this, we assigned an attractive potential between lipid head beads and the C-terminus beads of the TMD and LD to represent the charged residues. We also assigned an attractive potential among TMD hydrophobic beads and between TMD hydrophobic beads and lipid tail beads. Assignments of the attractive potential were based on the hydrophobicity of residues in the VAMP and Stx TMDs and residues in LDs near the C-terminus (52).

Both of these attractive potentials have the form of *V*_tt,attr_. We set a depth *ϵ* = 0.6 *k*_*B*_*T* and a decay range *w*_TMD,TMD_ = 1.6*D*_TMD_ for the attractive interaction among TMDs. The attraction potential parameters between TMD beads and lipid beads were set to prevent TMD pullout (see SI for details). We choose a depth *ϵ*_TMD,h_ = 1.2 *k*_*B*_*T* and *ϵ*_TMD,t_ = 1.8 *k*_*B*_*T* and a decay range *w*_TMD,i_ = 1.6*D*_*ij*_, where *D*_*ij*_ is the sum of the interacting bead radii.

All rigid bodies have the same mass *m* as lipid beads. Moment of inertia matrixes ***I*** for all rigid bodies were that of a uniform density cylinder with principal axis along the x-axis,

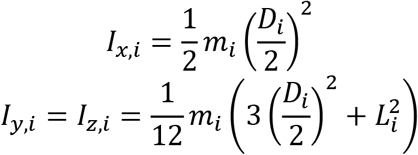

where i is either TMD or SNARE motif.

#### Molecular dynamics simulations

For simulations involving molecular fusogens, *N*_SNARE_ molecular fusogens are embedded into two synaptic vesicles with ~50 nm diameter (Fig. 1C). We used a rectangular box with sides *L*_x_ = *L*_y_ = 88 nm and *L*_z_ = 123 nm subject to periodic boundary conditions.

The vesicle tension is controlled by the osmotic pressure, which is created by placing solvent beads inside the lumen. Solvent beads interact only with lipid beads via the WCA potential and not with each other, similar to a previous membrane simulation study that used a phantom solvent to control system pressure (53). The concentration of solvent molecules in each vesicle was chosen to produce a desired pressure, which creates a tension in the vesicle membrane determined by the Young-Laplace law, *P*_solvent_ = 2*γ*_ves_/*R*_ves_, where *P*_solvent_ is the pressure produced by solvent beads, *γ*_ves_ is the vesicle membrane tension, and *R*_ves_ is the radius of the vesicle. The solvent beads have the same mass as lipid beads.

All MD simulations were run using the HOOMD-blue toolkit in the NVT ensemble using a Langevin thermostat (54–56). The simulation time step Δ*t* is set to 0.005*τ* (46–48, 57), where *τ* is the CG time unit. The translational drag was *γ*_*t*_ = *m*/*τ* for all beads (46–48) and rigid bodies. The rotational drag matrix was *γ*_rot *i,j*_ = *γ*_*t*_*δ*_*ij*_ for all rigid bodies.

### Brute force generates a large contact area and fuses vesicles inefficiently

The conceptually simplest way to fuse two membranes is to press them together. A previous simulation study induced fusion between two ~30 nm diameter CG lipid vesicles (43) by applying biasing forces to each bead of the lipid molecules. The biasing forces were released at the moment when lipid exchange between the two vesicles was initiated. How fusion is affected by the size of the vesicles and the magnitude of the applied force was not discussed.

Here, we used a similar procedure, varying the size of the vesicles as well the magnitude of the applied force. We simulated two vesicles of equal diameter 30 nm or 50nm with tension ~1 pN/nm. The applied force ranged from ~1460 pN to ~100 pN. The two types of vesicles were confined in a cylindrical wall of diameter 50 nm and 60 nm, respectively, to prevent them from sliding past each other. We then applied a constant force on every lipid bead in the two vesicles, pressing the vesicles towards each other. We found that this method successfully induced fusion between the two vesicles but required much larger forces on each vesicle than are provided by cellular fusogens.

In our simulations, pressing the two ~30 nm vesicle membranes together first resulted in the formation of a large flattened membrane contact zone with a radius ~10 nm (Fig. 2A). Multiple hemifusion connections were formed successively at the edge of this contact zone (Fig. 2B). The fate of these hemifusion connections depended on the magnitude of the applied force: if the applied force exceeded a critical force ~360 pN per vesicle, one of the hemifusion connections then expanded laterally along the contact zone edge and developed into a fusion pore (Fig. 2B). When the force was less than this value, hemifusion connections formed at the contact zone edge with lifetime ~1-10 μs and severed stochastically; no hemifusion connections were seen to develop into a fusion pore within ~1 ms, the total duration of each simulation. Pressing the two ~50 nm vesicles together, the fusion pathway was similar to that of the 30 nm vesicles, but the critical force and the contact zone radius increased to ~670 pN and ~20 nm, respectively. Interestingly, when the critical force, ~360 pN per 30nm diameter vesicle and ~670 pN per ~50 nm diameter vesicle, was applied, the contact zone area gives estimated pressures of ~10 atm and ~6 atm, respectively, agreeing with the reported minimum pressure to fuse two DMPC membranes measured using the surface force apparatus (58).

**Figure 2.**
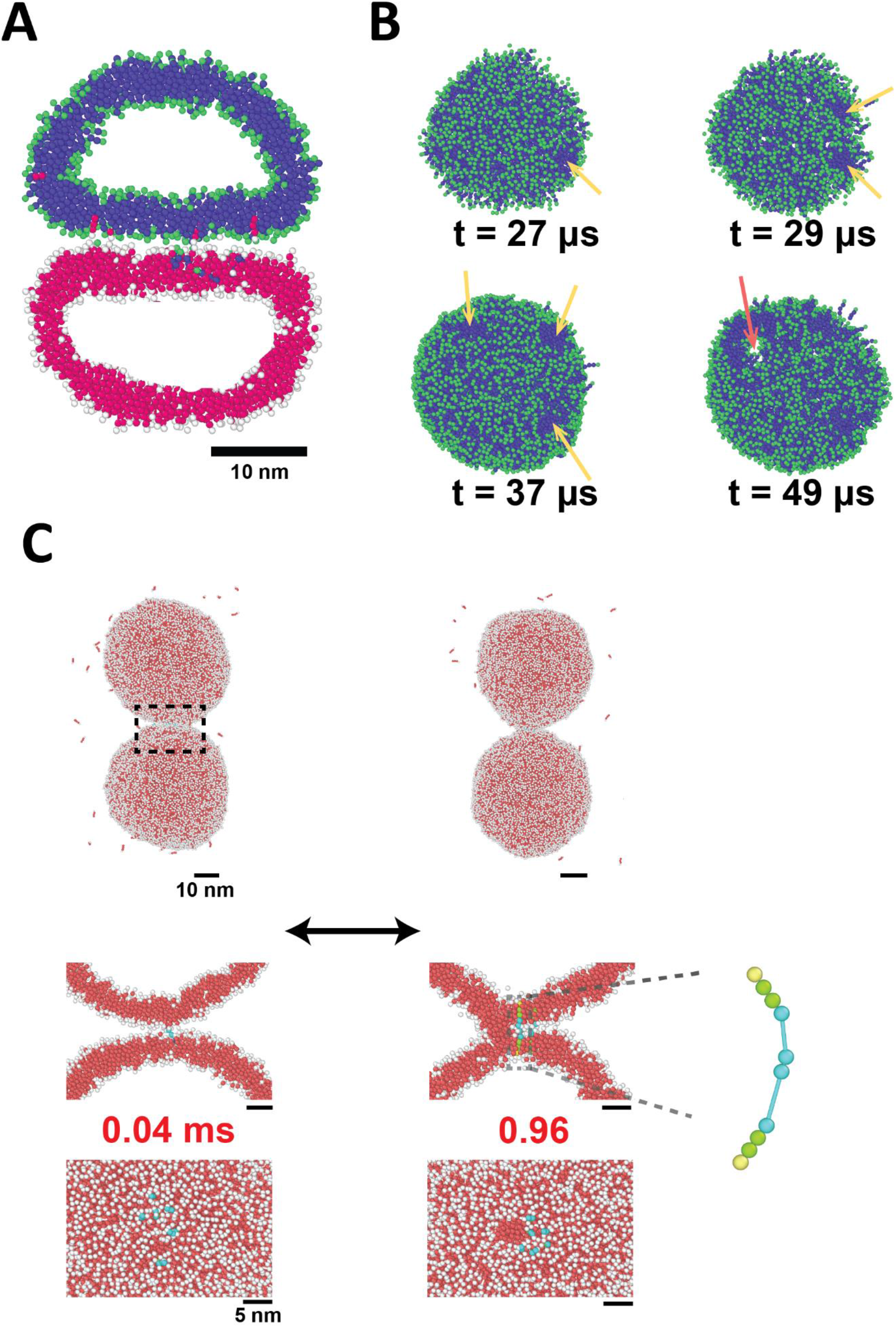
Brute force method mediating membrane fusion and the TMD-LD-TMD (TLT) fusogens. (A) Cross-section of two ~30 nm diameter vesicles being pressed together, showing the flattened membrane contact zone. Lipid head beads (green, white) and lipid tail beads (blue, magenta) are colored differently to distinguish the upper and lower vesicles. (B) Bottom view of a membrane patch at the membrane contact zone from the top vesicle shown in (A). First, a hemifusion connection formed at the edge of the contact zone, indicated by a yellow arrow. More hemifusion connections accumulated until a fusion pore developed in one of them (bottom right snapshot, orange arrow). (C) The TLT fusogen cannot fuse two vesicles together. The TLT fusogen consists of two TMDs and two LDs with the same color scheme as in Fig. 1D. Simulation snapshots of the whole system (top), the cross-section side view box region (middle), and the top view of the contact zone between vesicles (bottom) are shown. The fusogens cluster on the vesicles. TLT fusogens often lead to reversible hemifused stalk connections.

We note, however, that this brute force method is much less efficient than SNARE complexes at driving fusion because the total force required to mediate fusion between two ~30 nm vesicles, ~720 pN, is orders of magnitude larger than the force generated by SNAREpin zippering, ~18 pN (3). This is likely because the simple brute force approach generated a large membrane contact zone, weakening the pressure so that a larger force was needed to reach the ~10 atm fusion threshold (58).

### Non-rodlike membrane-bound fusogens produce transient hemifusion and no fusion

During cellular membrane fusion, the force pressing the membranes together is provided by a specialized molecular machinery interposed between the two fusing membranes. Thus we asked if fusogens providing a tight molecular connection between membranes were sufficient for fusion. We simulated a synthetic molecular fusogen complex (Fig. 2C), anchored in the two membranes with two TMDs, connected via two LDs, which we refer to as TMD-LD-TMD (TLT) fusogens. These are membrane-spanning complexes we imagine were produced by an earlier complexation event of two fusogens, one in each membrane. Unlike SNARE complexes, they are not rodlike. The LDs were represented by the bonded potential *V*_LD_(*r*) and are assumed to generate zippering forces of 18 pN, comparable to those of the neuronal SNARE complex (3).

We performed simulations to test whether such fusogens could mediate membrane fusion, and found that, although the TLT fusogens could sometimes produce hemifusion, they were unable to mediate membrane fusion within the ~1 ms simulation time because the driving force pressing the membranes together was lost after hemifusion was achieved. We simulated two ~ 50 nm vesicles connected via six copies of the TLT fusogens. In the simulations, tension in the LDs caused the TMDs to aggregate at the point of closest approach between the two membranes (Fig. 2C). In most simulations, hemifusion was not achieved during the 1 ms run (*n* = 12 of 20 simulations). In the remaining runs, a hemifusion connection (1-30 μs) formed near the cluster of fusogens (Fig. 2C) and then stochastically severed, similar to simulations using the brute force method (*n* = 8 of 20 runs). During these brief hemifused episodes, the LDs connecting the two TMDs were able to contract to a minimal length by migrating to the edge of the hemifused zone, fig. 2C. Thus, the driving force for fusion was lost following hemifusion, and the TLT fusogens were unable to drive full fusion. The vesicles were in a hemifused state for only a tiny fraction of the time; during an entire ~1 ms simulation, only 0-2 hemifusion episodes typically occurred, lasting ~ 1-30 μs.

### SNARE-like molecular fusogens clear the contact zone, drive long-lived hemifusion diaphragms, and drive the transition from hemifusion to fusion

The core of the cellular fusion machinery, the SNARE complex, is a ~ 2 nm thick and ~ 10 nm long rod-shaped coiled-coil. Viral fusion proteins such as the spike protein of SARS-CoV-2 (59) and hemagglutinin of influenza (60) are also rodlike structures in the folded state. Our previous simulation studies of SNARE-mediated neurotransmitter release suggested that the rodlike structure of the SNARE complexes gave rise to entropic forces that organize the SNAREs into a ring-like arrangement and drive expansion of the ring, pressing the membranes together for fusion (33, 34). We therefore hypothesized that relative to the simple transmembrane TLT complex, a rodlike structure might stabilize hemifusion and facilitate the transition from hemifusion to fusion.

We performed simulations using model SNARE-like fusogens in which the coiled-coil of the SNARE complex was represented with a rigid rod-shaped domain comprising ten beads, each 2 nm in diameter (Fig. 1D). The SNARE-like fusogens spontaneously cleared themselves from the contact zone, and organized into a ring. SNARE-like fusogens successfully drove full fusion of the two vesicles in the majority of simulations.

Unlike the TLT fusogens, the SNARE-like fusogens did not cluster at the point of closest approach between the two membranes. Initially, 6 SNARE-like fusogens were placed into a ring with the coiled-coil domains pointing outward with a radius of 10 nm connecting the two ~50 nm diameter vesicles (Fig. 1C). The fusogens maintained this organization, though the radius of the ring decreased to 5.7 ± 0.8 nm (Fig. 3). The fusogens spontaneously vacated the contact zone between the vesicles due to collisions between the long rods and between the rods and the membrane. These collision forces are of entropic origin, tending to expand the ring of SNARE-like fusogens: larger rings are less crowded and give the fusogens greater orientational entropy (33, 34).

**Figure 3.**
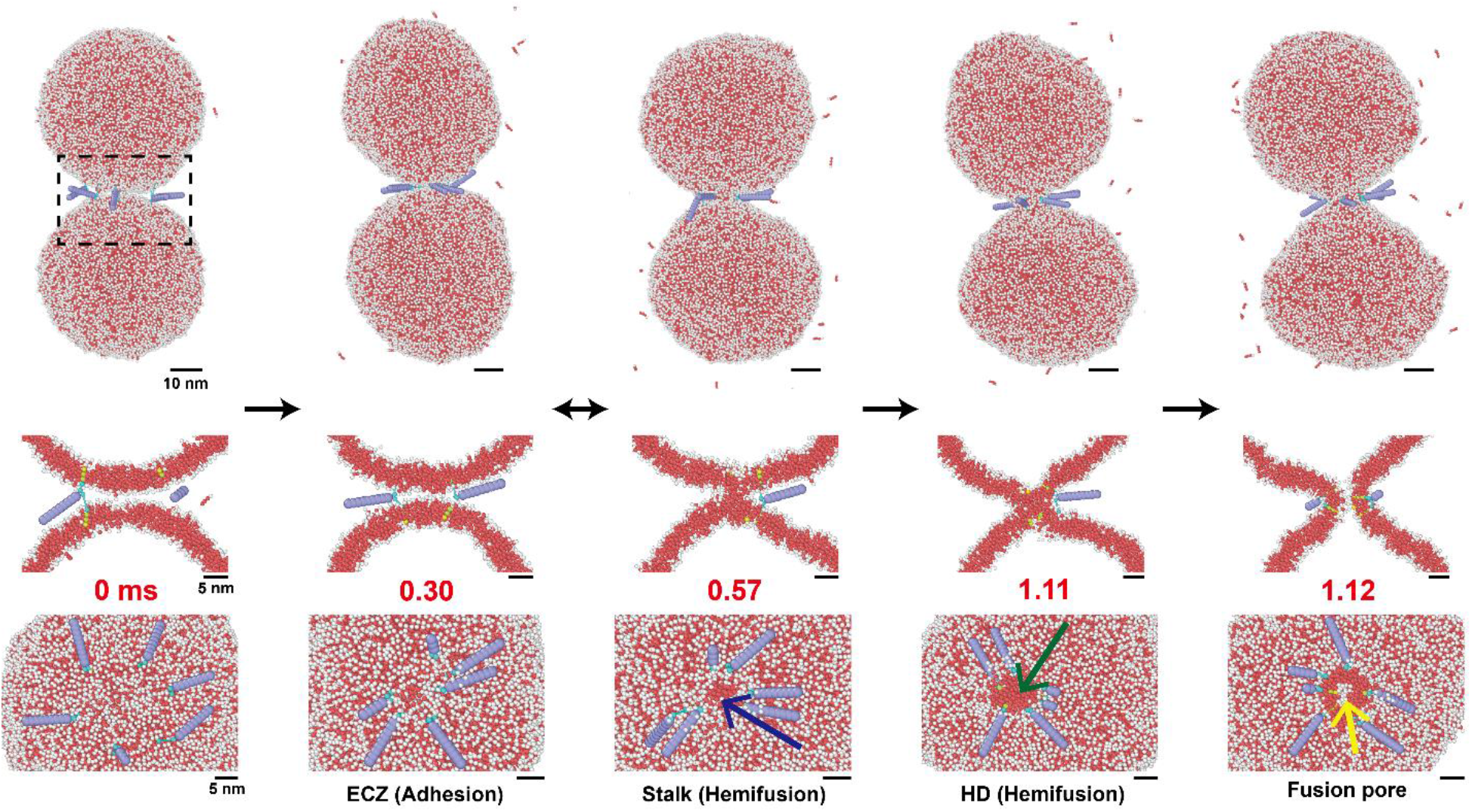
Snapshots showing the SNARE-mediated membrane fusion pathway. Top: Side view of fusing vesicles. Scale bar: 10 nm. Middle: Cross-section of the box region in the system. Scale bar: 5 nm. Bottom: Top view of the SNAREpin ring, including the bottom vesicle membrane. Scale bar: 5 nm. The membrane adhesion zone develops early, after 1 μs.

These entropic forces pushed the fusogens to the edge of the contact zone (Fig. 3), where the SNARE-like fusogens were able to successfully drive the opening of a fusion pore. In *n* = 35 out of 40 runs, a hemifusion connection formed at the edge of the contact zone (Fig. 3). Once the hemifusion connection was formed, the fusogens migrated radially inward toward the edge of the hemifusion connection, driven by the zippering force. In *n* = 23 simulations, the hemifusion connection expanded into a hemifusion diaphragm (Fig. 3), in which a simple pore then developed, yielding fusion (Fig. 3). Note that, unlike the TLT constructs, the SNARE-like fusogens stabilized hemifusion connections which were sometimes long-lived (reach ~ 0.1 ms) via entropic force.

### Membrane fusion requires a sufficiently long SNARE motif to generate sufficient entropic force

So far, our simulations have shown that rod-shaped SNARE-like fusogens successfully drive membrane fusion, while non-rodlike TLT fusogens do not. We reasoned that this is due to entropic forces associated with the rod-shaped coiled-coil domain that tend to push the SNARE-like fusogens radially outward. To test this hypothesis, we performed simulations varying the length of the coiled-coil domain in the SNARE-like fusogens. A short coiled-coil domain would be expected to produce little to no entropic force and would therefore be expected to drive transient hemifusion, but not fusion.

We modeled SNARE “length mutants” with a SNARE motif lengths 2 nm, 6 nm, and 14 nm, compared to the “wildtype” length of 10 nm (Fig. 4A). We performed simulations using 6 copies of each mutant (*n* = 20) and measured the fraction of simulations producing hemifusion or hemifusion followed by fusion.

**Figure 4.**
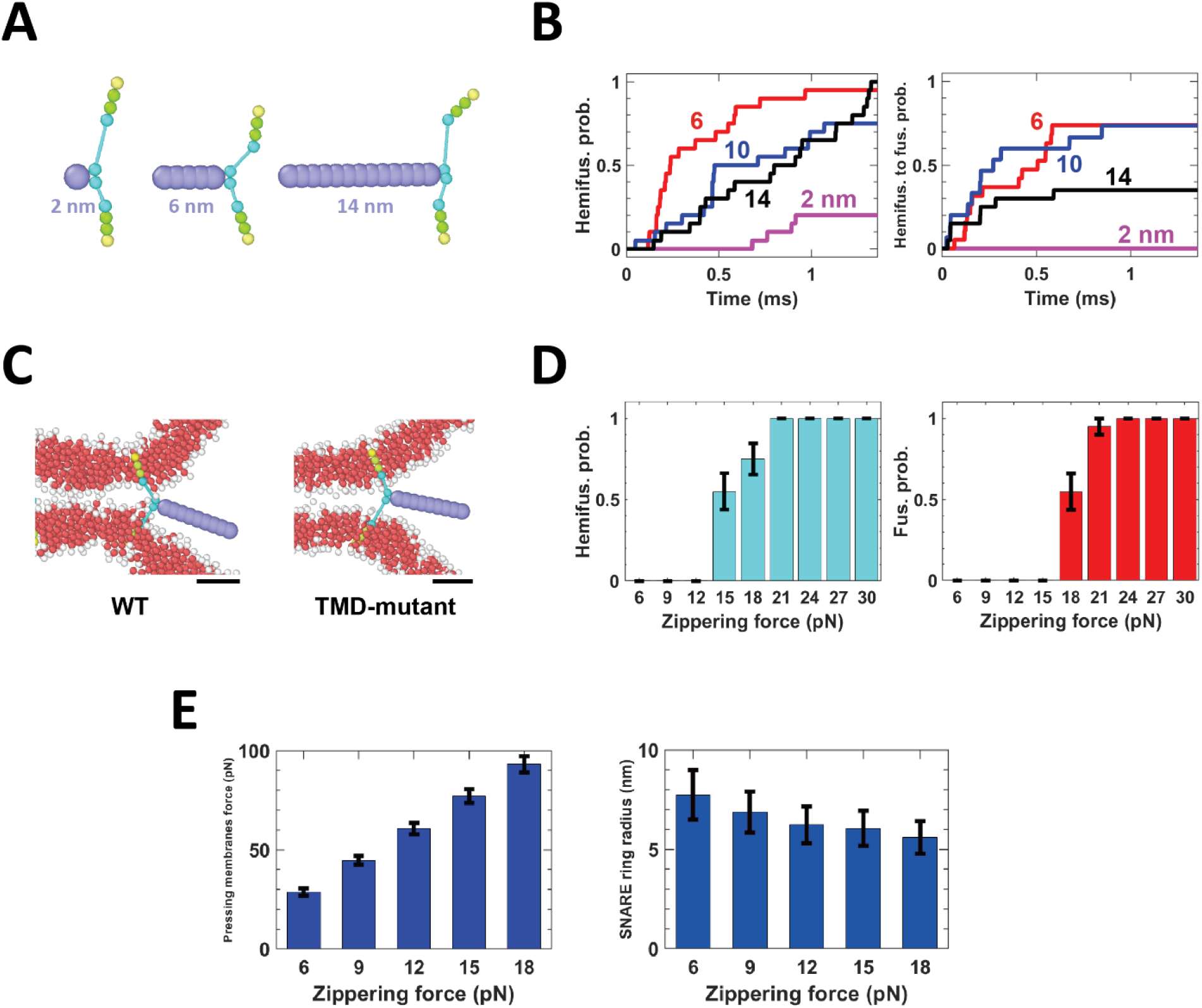
SNARE mutants affect membrane fusion kinetics. (A) Two SNARE mutants with 2 nm (left), 6 nm (middle), and 14 nm (right) SNARE motif. (B) Cumulative of the fraction of simulations (n=20) achieved hemifusions (left) and fusion after hemifusion happened (right) using mutants with different SNARE motif lengths (6 SNAREs) during 1.36 ms simulation time. For more details, please see Fig. S1. (C) Simulation snapshot of the WT SNARE with local membrane thinning (left) and SNARE mutants with neutrally charged TMD C-terminus and LD C-terminus residues to switch off local thinning (right). Scale bar: 5 nm. (D) Fraction of 6-SNARE-mediated fusion simulations (n=20) resulted in hemifusion (left) and fusion (right) under each SNARE zippering force within 1.36 ms. (E) Force pressing membrane together (left) and the SNARE ring radius (right) under each value of zippering force before hemifusion happened. Measurement was conducted every ~10 μs. Error bars: SD.

Simulations showed that stable hemifusion and fusion required coiled-coil domains longer than 2 nm, consistent with the conclusion that entropic forces expanding the ring of fusogens stabilized hemifusion and drove the transition from hemifusion to fusion. With a coiled-coil length of 2 nm, fusion occurred in zero runs, while with a longer coiled-coil, fusion occurred after a certain waiting time (Fig. 4B). Similar to the TLT fusogens, SNARE-like fusogens with a 2 nm coiled-coil domain drove transient hemifusion in *n* = 4 runs (out of 20) (Fig. 4B).

### TMD-induced membrane thinning facilitates fusion

Mathematical models and simulations have shown that membrane insertions and TMDs can locally perturb the membrane thickness (28, 61). This effect was proposed to explain the spontaneous sorting of Syntaxin TMDs into membrane domains of various compositions (35). In VAMP2 and Syntaxin-1, several residues at the interface between the LD and TMD are positively charged, and the TMDs have a negatively charged carboxyl group at their C-terminus. These charged groups strongly prefer to interact with polar lipid head groups. Since the length of the TMDs (~3 nm) is shorter than the membrane thickness (~5 nm), TMDs can thin the membrane locally. The locally thinned membrane has a shape of a dimple with positive curvature at its edge. These positive curvature regions may facilitate membrane fusion by exposing the hydrophobic membrane interiors toward the opposing membrane, enhancing lipid exchange (62), and inducing membrane stress that may be relaxed by fusion.

Interestingly, in simulations, the TMDs of the SNARE-like fusogens thinned the membrane locally and formed dimples on each membrane (Fig. 4C) due to the mismatch in the lengths of the hydrophobic regions of the membrane and the SNARE TMDs. We hypothesized that the membrane stress induced by the hydrophobic mismatch in our simulations might promote membrane fusion.

We therefore performed simulations using SNARE-like fusogens lacking attractive interactions between the N and C terminal beads of the TMDs and the lipid head beads. This mutation abolished the ability of the fusogens to induce local thinning of the membrane (Fig. 4C). Simulations used 6 fusogens.

In these simulations with abolished membrane thinning, no fusion occurred during the ~ 1ms running time (*n* = 10 runs). To quantify the decrease in thinning, we measured the membrane insertion depths of the TMD C-terminus bead and the LD C-terminus bead for both the SNARE mutants and the WT SNAREs throughout the simulations. For the WT SNARE, the two beads were significantly closer to surrounding lipid head beads than the SNARE mutant (Fig. 4C), indicating that the WT SNAREs could thin the membrane much more potent.

### Membrane fusion requires a sufficiently strong zippering force

Optical tweezer experiments have shown that a single neuronal SNARE complex can provide a zippering force of ~ 18 pN (3). This finding has been reproduced using CG simulations (27). Based on our simulations showing that membrane fusion requires the membranes to be pressed together with force greater than a critical value, we hypothesized that membrane fusion requires a sufficiently large zippering force.

We therefore performed simulations using 6 SNARE-like fusogens, varying the magnitude of the zippering force from 6 pN to 18 pN. A zippering force of 18 pN per SNARE or more was required for fusion, while 15 pN was required for hemifusion (Fig. 4D). Simulations with a zippering force less than 18 pN produced no fusion; the fraction of simulations in which fusion occurred increased as the zippering force increased, reaching 100% with a zippering force of 24 pN. Membrane fusion was faster as the zippering force increased beyond the 18 pN requirement (Fig. 4D).

We measured the radius of the ring of SNAREs that formed prior to hemifusion (fig. 4D) and the total force pressing the two membranes together (Fig. 4E). Interestingly, we observed a smaller SNARE ring with a larger SNARE zippering force. With a larger zippering force, the SNAREs were more spatially confined to generate higher entropic force to balance the in-plane component of the zippering force. When the zippering force reached 15 pN, the pressure acting on the vesicle-vesicle contact zone within the snare ring reached ~10 atm, comparable to the value required in brute force simulations, suggesting a minimum zippering force of this magnitude should be expected. However, we note that the exact value of the minimum zippering force may be sensitive to the parameters of our model and the duration of our simulations.

### Membrane fusion is faster with more SNAREs

Whether fusion requires a minimum number of SNARE complexes is controversial, and reports of the number required range widely from 1 to 10 (5, 7–9, 11–13, 32). In our previous simulation studies, we found that no requirement for the number of SNAREs, but fusion was faster with more SNAREs (33, 34). Here, we performed simulations using varying numbers of SNARE-like fusogens to investigate the dependence of fusion kinetics on the number of SNAREs at the fusion site.

Consistent with previous simulation studies, fusion was faster with more SNARE-like fusogens. With more than 3 fusogens, the fraction of runs in which fusion occurred increased with additional SNARE-like fusogens, while the fraction of runs in which hemifusion occurred was approximately unchanged (Fig. 5A). Similarly, the delay from hemifusion to fusion deceased with additional fusogens. Our results suggest that the two membranes are more likely to be trapped in the hemifusied state with fewer SNAREs (“dead-end hemifusion”).

**Figure 5.**
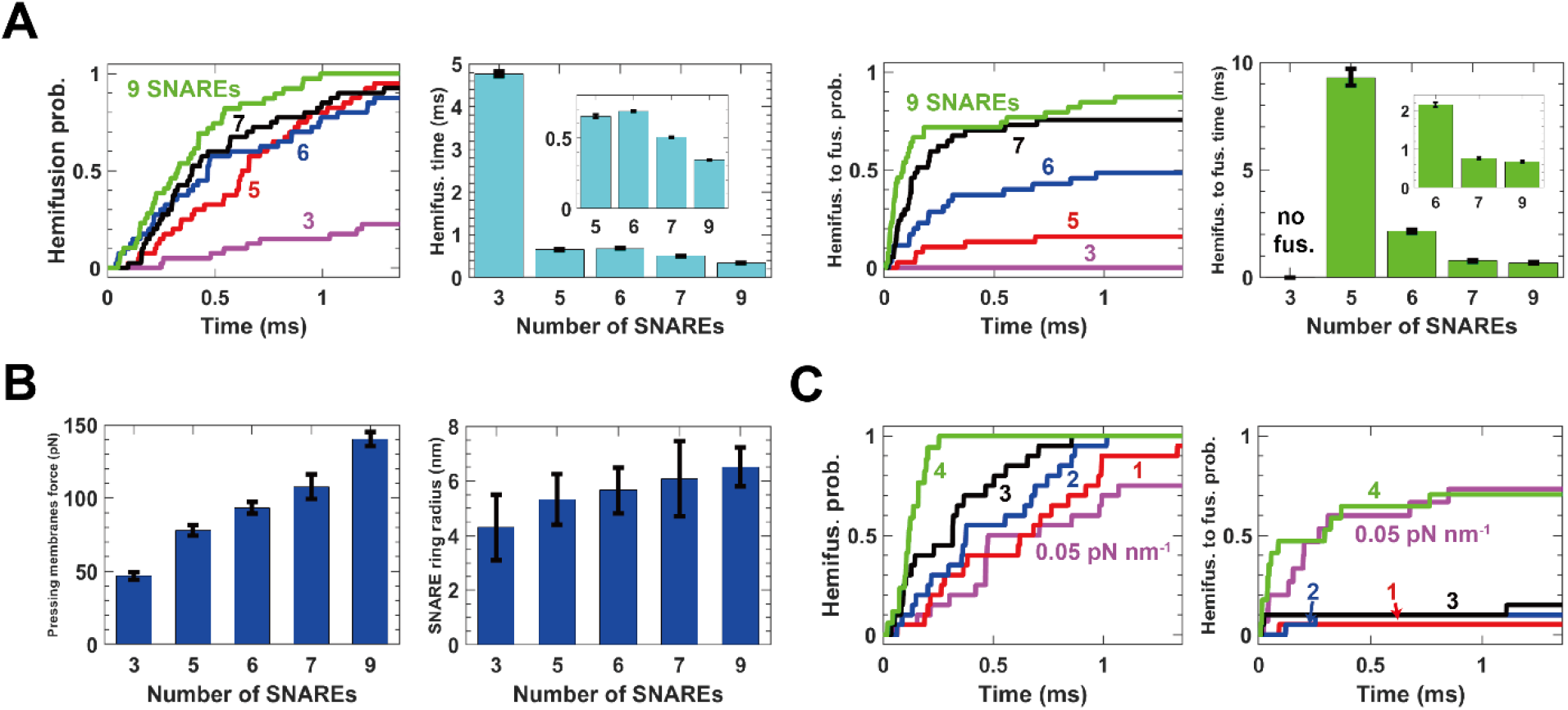
Effect of number of SNAREs and membrane tension on the hemifusion and fusion kinetics. (A) Cumulative distribution function describing the fraction of simulations (n=40) achieving hemifusion and fusion following hemifusion, for different numbers of SNAREs during the 1.36 ms simulation time. Mean hemifusion and fusion times are shown. The insets are the zoomed in view for hemifusion time of 5-9 SNAREs and hemifusion to fusion time for 6-9 SNAREs, respectively. Error bars: SDs of mean time. For details, see Fig. S2. (B) Force pressing membrane together and the SNARE ring radius (right) under each value of zippering force before hemifusion happened. Measurement was conducted every ~10 μs. Error bars: SD. (C) The cumulative distribution function of the fraction of 6-SNARE-mediated simulations (n=20) resulted in hemifusion (left) and fusion after hemifusion happened (right) under each membrane tension during 1.36 ms simulation time. For more details, please see Fig. S3.

With fewer than 3 fusogens, we observed no fusion events over the course of simulations lasting ~1 ms (*n* = 40). We speculate that this apparent requirement for 3 or more fusogens could be caused by our limited simulation time. Indeed, extrapolating the membrane fusion time to fewer than 3 SNARE-like fusogens produced estimates in excess of the total duration of our simulations.

Two mechanisms influence the fusion kinetics as the number of SNAREs changes. Firstly, the total zippering force provided by all SNAREs increases with more SNAREs present, resulting in a greater force pressing the two membranes together (Fig. 5B). In the previous section, we showed that the fusion rate was faster with a larger zippering force per SNARE-like fusogen; thus, a greater net pressing force is expected to speed fusion. Secondly, the entropic forces expanding the SNARE ring are enhanced with additional SNAREs (Fig. 5B). Thus, with fewer SNAREs, a weaker force presses the membranes together, reducing fusion rates.

### Membrane tension increases hemifusion and fusion rates

Experimental studies suggested that membrane tension activated exocytosis during the migration of fibroblasts (63, 64). In an in vitro study of SNARE-mediated fusion lipid mixing between large unilamellar vesicles (LUVs) and either giant unilamellar vesicles (GUVs) or planar membranes increased sharply when the tension of the GUV or planar membrane was elevated beyond ~3-4 pN/nm (65), much higher than typical cellular membrane tensions, ~0.04 pN/nm (66). MD simulations of protein-free fusion also found that fusion required high tensions (67); however, the reported threshold value required for fusion of ~50 pN/nm is several times greater than typical membrane rupture tensions ~5-10 pN/nm, possibly undetected due to the ~1μs timescales of their simulations. It has been hypothesized that membrane tension facilitates membrane fusion directly by lowering the energy barrier to fusion (68). Thus we performed simulations to investigate the possible effects of membrane tension on SNARE-mediated membrane fusion kinetics.

We used 6 SNARE-like fusogens and performed simulations under different membrane tension ranging from 0.05 pN/nm to 4 pN/nm (*n* = 20 runs). Membrane tension was set by placing solvent beads in each vesicle with a concentration chosen to produce the desired tension based on the Young-Laplace law and the ideal gas law. Solvent beads interact only with lipids and are thus expected to be well-described by the ideal gas law. We found that hemifusion events occurred more frequently at higher membrane tensions (Fig. 5C), and fusion was delayed as tension increases from 0.05 pN/nm, but fusion rate was restored when the tension reached the rupture tension in our simulation, 4 pN/nm.

## Conclusions

Here we used 4-bead lipids interacting via the well-tested Cooke-Deserno force field (46–48) to build DPPC vesicles. We controlled the vesicle membrane tension with a fictitious intravesicle gas, and used SNAREpin-like fusogens whose zippering was mimicked with an 18 pN zippering force as measured experimentally (3) (Fig. 1). With this UCG system, the UCG SNAREpins fused 50-nm vesicles on ms timescales along a reproducible fusion pathway: SNARE-SNARE and SNARE-membrane entropic forces cleared the fusion site and pressed the vesicles into an extended contact zone (ECZ), drove a stalk to nucleate at the high curvature ECZ boundary, and drove stalk expansion into a long-lived hemifusion diaphragm (HD) in which a simple pore completed fusion.

Consistent with enhanced neurotransmitter release rates observed with cells with mutations so that more SNAREpins are at the fusion site (32, 69), in simulations more SNARES generated higher entropic forces and fused membranes faster (34), from ~ 8.9 ms (5 SNAREs) to ~ 0.7 ms (9 SNAREs). TMD-mediated membrane thinning (35, 36) and zippering forces promoted fusion. In simulations with short fusogens, fusion and entropic forces were abolished. In summary, our results suggest that the functional significance of the rodlike SNAREpin shape is for fusogenicity, endowing SNARE complexes with the ability to exert entropic forces that drive hemifusion and fusion on ms timescales.

## Acknowledgements

Research reported in this publication was supported by the National Institute of General Medical Sciences of the National Institutes of Health under award number R01GM117046 to B.O. The content is solely the responsibility of the authors and does not necessarily represent the official views of the National Institutes of Health. We acknowledge computing resources from Columbia University’s Shared Research Computing Facility project, which is supported by NIH Research Facility Improvement Grant 1G20RR030893-01, and associated funds from the New York State Empire State Development, Division of Science Technology and Innovation (NYSTAR) Contract C090171, both awarded April 15, 2010.

## References

1. T. C. Sudhof, J. E. Rothman, Membrane fusion: grappling with SNARE and SM proteins. Science 323, 474–477 (2009).

2. T. Wang, L. Li, W. Hong, SNARE proteins in membrane trafficking. Traffic 18, 767–775 (2017).

3. Y. Gao et al., Single reconstituted neuronal SNARE complexes zipper in three distinct stages. Science 337, 1340–1343 (2012).

4. L. Ma et al., alpha-SNAP Enhances SNARE Zippering by Stabilizing the SNARE Four-Helix Bundle. Cell Rep 15, 531–539 (2016).

5. Y. Hua, R. H. Scheller, Three SNARE complexes cooperate to mediate membrane fusion. Proc Natl Acad Sci U S A 98, 8065–8070 (2001).

6. X. Han, C. T. Wang, J. Bai, E. R. Chapman, M. B. Jackson, Transmembrane segments of syntaxin line the fusion pore of Ca2+-triggered exocytosis. Science 304, 289–292 (2004).

7. R. Sinha, S. Ahmed, R. Jahn, J. Klingauf, Two synaptobrevin molecules are sufficient for vesicle fusion in central nervous system synapses. Proc Natl Acad Sci U S A 108, 14318–14323 (2011).

8. R. Mohrmann, H. de Wit, M. Verhage, E. Neher, J. B. Sorensen, Fast vesicle fusion in living cells requires at least three SNARE complexes. Science 330, 502–505 (2010).

9. G. van den Bogaart et al., One SNARE complex is sufficient for membrane fusion. Nat Struct Mol Biol 17, 358–364 (2010).

10. L. Shi et al., SNARE proteins: one to fuse and three to keep the nascent fusion pore open. Science 335, 1355–1359 (2012).

11. E. Karatekin et al., A fast, single-vesicle fusion assay mimics physiological SNARE requirements. Proc Natl Acad Sci U S A 107, 3517–3521 (2010).

12. M. K. Domanska, V. Kiessling, A. Stein, D. Fasshauer, L. K. Tamm, Single vesicle millisecond fusion kinetics reveals number of SNARE complexes optimal for fast SNARE-mediated membrane fusion. J Biol Chem 284, 32158–32166 (2009).

13. J. M. Hernandez, A. J. Kreutzberger, V. Kiessling, L. K. Tamm, R. Jahn, Variable cooperativity in SNARE-mediated membrane fusion. Proc Natl Acad Sci U S A 111, 12037–12042 (2014).

14. R. Jahn, R. H. Scheller, SNAREs--engines for membrane fusion. Nat Rev Mol Cell Biol 7, 631–643 (2006).

15. R. Jahn, D. Fasshauer, Molecular machines governing exocytosis of synaptic vesicles. Nature 490, 201–207 (2012).

16. V. Knecht, H. Grubmuller, Mechanical coupling via the membrane fusion SNARE protein syntaxin 1A: a molecular dynamics study. Biophys J 84, 1527–1547 (2003).

17. B. Liang, D. Dawidowski, J. F. Ellena, L. K. Tamm, D. S. Cafiso, The SNARE motif of synaptobrevin exhibits an aqueous-interfacial partitioning that is modulated by membrane curvature. Biochemistry 53, 1485–1494 (2014).

18. C. S. Kim, D. H. Kweon, Y. K. Shin, Membrane topologies of neuronal SNARE folding intermediates. Biochemistry 41, 10928–10933 (2002).

19. N. A. Lakomek, H. Yavuz, R. Jahn, A. Perez-Lara, Structural dynamics and transient lipid binding of synaptobrevin-2 tune SNARE assembly and membrane fusion. Proc Natl Acad Sci U S A 116, 8699–8708 (2019).

20. B. Liang, V. Kiessling, L. K. Tamm, Prefusion structure of syntaxin-1A suggests pathway for folding into neuronal trans-SNARE complex fusion intermediate. Proc Natl Acad Sci U S A 110, 19384–19389 (2013).

21. J. F. Ellena et al., Dynamic structure of lipid-bound synaptobrevin suggests a nucleation-propagation mechanism for trans-SNARE complex formation. Proc Natl Acad Sci U S A 106, 20306–20311 (2009).

22. J. A. McNew, T. Weber, D. M. Engelman, T. H. Sollner, J. E. Rothman, The length of the flexible SNAREpin juxtamembrane region is a critical determinant of SNARE-dependent fusion. Mol Cell 4, 415–421 (1999).

23. P. Zhou, T. Bacaj, X. Yang, Z. P. Pang, T. C. Sudhof, Lipid-anchored SNAREs lacking transmembrane regions fully support membrane fusion during neurotransmitter release. Neuron 80, 470–483 (2013).

24. J. Kesavan, M. Borisovska, D. Bruns, v-SNARE actions during Ca(2+)-triggered exocytosis. Cell 131, 351–363 (2007).

25. Y. Hu, L. Zhu, C. Ma, Structural Roles for the Juxtamembrane Linker Region and Transmembrane Region of Synaptobrevin 2 in Membrane Fusion. Front Cell Dev Biol 8, 609708 (2020).

26. M. P. Durrieu, R. Lavery, M. Baaden, Interactions between neuronal fusion proteins explored by molecular dynamics. Biophys J 94, 3436–3446 (2008).

27. S. Sharma, M. Lindau, Molecular mechanism of fusion pore formation driven by the neuronal SNARE complex. Proc Natl Acad Sci U S A 115, 12751–12756 (2018).

28. H. J. Risselada, C. Kutzner, H. Grubmuller, Caught in the act: visualization of SNARE-mediated fusion events in molecular detail. Chembiochem 12, 1049–1055 (2011).

29. R. Schneggenburger, E. Neher, Intracellular calcium dependence of transmitter release rates at a fast central synapse. Nature 406, 889–893 (2000).

30. T. Sakaba, A. Stein, R. Jahn, E. Neher, Distinct kinetic changes in neurotransmitter release after SNARE protein cleavage. Science 309, 491–494 (2005).

31. L. Y. Wang, E. Neher, H. Taschenberger, Synaptic vesicles in mature calyx of Held synapses sense higher nanodomain calcium concentrations during action potential-evoked glutamate release. J Neurosci 28, 14450–14458 (2008).

32. C. Acuna et al., Microsecond Dissection of Neurotransmitter Release: SNARE-Complex Assembly Dictates Speed and Ca2+ Sensitivity. Neuron 82, 1088–1100 (2014).

33. H. Mostafavi et al., Entropic forces drive self-organization and membrane fusion by SNARE proteins. Proc Natl Acad Sci U S A 114, 5455–5460 (2017).

34. Z. A. McDargh, A. Polley, B. O’Shaughnessy, SNARE-mediated membrane fusion is a two-stage process driven by entropic forces. FEBS Lett 592, 3504–3515 (2018).

35. D. Milovanovic et al., Hydrophobic mismatch sorts SNARE proteins into distinct membrane domains. Nat Commun 6, 5984 (2015).

36. Y. G. Smirnova, H. J. Risselada, M. Muller, Thermodynamically reversible paths of the first fusion intermediate reveal an important role for membrane anchors of fusion proteins. Proc Natl Acad Sci U S A 116, 2571–2576 (2019).

37. J. Nikolaus, M. Stockl, D. Langosch, R. Volkmer, A. Herrmann, Direct visualization of large and protein-free hemifusion diaphragms. Biophys J 98, 1192–1199 (2010).

38. J. M. Warner, B. O’Shaughnessy, The hemifused state on the pathway to membrane fusion. Phys Rev Lett 108, 178101 (2012).

39. J. M. Warner, B. O’Shaughnessy, Evolution of the hemifused intermediate on the pathway to membrane fusion. Biophys J 103, 689–701 (2012).

40. J. M. Hernandez et al., Membrane fusion intermediates via directional and full assembly of the SNARE complex. Science 336, 1581–1584 (2012).

41. J. Diao et al., Synaptic proteins promote calcium-triggered fast transition from point contact to full fusion. Elife 1, e00109 (2012).

42. W. D. Zhao et al., Hemi-fused structure mediates and controls fusion and fission in live cells. Nature 534, 548–552 (2016).

43. M. J. Stevens, J. H. Hoh, T. B. Woolf, Insights into the molecular mechanism of membrane fusion from simulation: Evidence for the association of splayed tails. Physical Review Letters 91, 188102 (2003).

44. B. L. Sabatini, W. G. Regehr, Timing of neurotransmission at fast synapses in the mammalian brain. Nature 384, 170–172 (1996).

45. B. L. Sabatini, W. G. Regehr, Timing of synaptic transmission. Annu Rev Physiol 61, 521–542 (1999).

46. I. R. Cooke, M. Deserno, Solvent-free model for self-assembling fluid bilayer membranes: stabilization of the fluid phase based on broad attractive tail potentials. J Chem Phys 123, 224710 (2005).

47. I. R. Cooke, K. Kremer, M. Deserno, Tunable generic model for fluid bilayer membranes. Phys Rev E Stat Nonlin Soft Matter Phys 72, 011506 (2005).

48. G. Illya, M. Deserno, Coarse-grained simulation studies of peptide-induced pore formation. Biophys J 95, 4163–4173 (2008).

49. B. A. Lewis, D. M. Engelman, Lipid bilayer thickness varies linearly with acyl chain length in fluid phosphatidylcholine vesicles. J Mol Biol 166, 211–217 (1983).

50. A. Filippov, G. Oradd, G. Lindblom, Lipid lateral diffusion in ordered and disordered phases in raft mixtures. Biophys J 86, 891–896 (2004).

51. R. B. Sutton, D. Fasshauer, R. Jahn, A. T. Brunger, Crystal structure of a SNARE complex involved in synaptic exocytosis at 2.4 angstrom resolution. Nature 395, 347–353 (1998).

52. A. Stein, G. Weber, M. C. Wahl, R. Jahn, Helical extension of the neuronal SNARE complex into the membrane. Nature 460, 525–528 (2009).

53. O. Lenz, F. Schmid, A simple computer model for liquid lipid bilayers. Journal of Molecular Liquids 117, 147–152 (2005).

54. J. A. Anderson, J. Glaser, S. C. Glotzer, HOOMD-blue: A Python package for high-performance molecular dynamics and hard particle Monte Carlo simulations. Computational Materials Science 173, 109363 (2020).

55. T. D. Nguyen, C. L. Phillips, J. A. Anderson, S. C. Glotzer, Rigid body constraints realized in massively-parallel molecular dynamics on graphics processing units. Computer Physics Communications 182, 2307–2313 (2011).

56. J. Glaser, X. Zha, J. A. Anderson, S. C. Glotzer, A. Travesset, Pressure in rigid body molecular dynamics. Computational Materials Science 173, 109430 (2020).

57. S. Foley, M. Deserno, Stabilizing Leaflet Asymmetry under Differential Stress in a Highly Coarse-Grained Lipid Membrane Model. J Chem Theory Comput 16, 7195–7206 (2020).

58. J. Y. Wong, C. K. Park, M. Seitz, J. Israelachvili, Polymer-cushioned bilayers. II. An investigation of interaction forces and fusion using the surface forces apparatus. Biophysical Journal 77, 1458–1468 (1999).

59. Y. Cai et al., Distinct conformational states of SARS-CoV-2 spike protein. Science 369, 1586–1592 (2020).

60. P. A. Bullough, F. M. Hughson, J. J. Skehel, D. C. Wiley, Structure of influenza haemagglutinin at the pH of membrane fusion. Nature 371, 37–43 (1994).

61. H. Aranda-Espinoza, A. Berman, N. Dan, P. Pincus, S. J. B. j. Safran, Interaction between inclusions embedded in membranes. 71, 648–656 (1996).

62. H. T. McMahon, M. M. Kozlov, S. Martens, Membrane curvature in synaptic vesicle fusion and beyond. Cell 140, 601–605 (2010).

63. N. C. Gauthier, M. A. Fardin, P. Roca-Cusachs, M. P. Sheetz, Temporary increase in plasma membrane tension coordinates the activation of exocytosis and contraction during cell spreading. Proc Natl Acad Sci U S A 108, 14467–14472 (2011).

64. N. C. Gauthier, O. M. Rossier, A. Mathur, J. C. Hone, M. P. Sheetz, Plasma membrane area increases with spread area by exocytosis of a GPI-anchored protein compartment. Mol Biol Cell 20, 3261–3272 (2009).

65. T. T. Kliesch et al., Membrane tension increases fusion efficiency of model membranes in the presence of SNAREs. Sci Rep 7, 12070 (2017).

66. J. Y. Tinevez et al., Role of cortical tension in bleb growth. Proc Natl Acad Sci U S A 106, 18581–18586 (2009).

67. J. C. Shillcock, R. Lipowsky, Tension-induced fusion of bilayer membranes and vesicles. Nat Mater 4, 225–228 (2005).

68. N. C. Gauthier, T. A. Masters, M. P. Sheetz, Mechanical feedback between membrane tension and dynamics. Trends Cell Biol 22, 527–535 (2012).

69. S. H. Gerber et al., Conformational switch of syntaxin-1 controls synaptic vesicle fusion. Science 321, 1507–1510 (2008).

