## Supporting Information for "Why are SNAREpins rod-shaped?"

### Supplementary methods

**Coarse-grained lipid representation developed in ref. (1-3)**
We use the force fields developed in refs. (1-3). Each lipid molecule is represented by 4 beads; we use the latter here. A lipid is a linear sequence of 1 head (h) bead followed by 3 tail (t) beads. Bonds exist between each successive pair of beads and between the first and the last bead. Potentials of bonded and non-bonded interactions are given below, and the parameters are in Table S1.

**Non-bonded interactions**
Beads interact via a Weeks-Chandler-Andersen potential (WCA),

$$V_{\mathrm{ij,wca}}(r_{\mathrm{ij}}) = \begin{cases} 4\epsilon\left[\left(\frac{b_{\mathrm{ij}}}{r_{\mathrm{ij}}}\right)^{12} - \left(\frac{b_{\mathrm{ij}}}{r_{\mathrm{ij}}}\right)^{6} + \frac{1}{4}\right], & r \le r_c \\ 0, & r > r_c \end{cases}$$

where i and j are either h or t. An attractive potential $V_{\mathrm{tt,attr}}$ represents the hydrophobic effect among all tail beads,

$$V_{\mathrm{tt,attr}}(r_{\mathrm{tt}}) = \begin{cases} -\epsilon, & r < r_c \\ -\epsilon\cos^2\frac{\pi(r-r_c)}{2w_c}, & r_c \le r \le r_c + w_c. \\ 0, & r > r_c + w_c \end{cases}$$

The attractive well has a depth of $\epsilon$ and a decay range $w_c$. $b_{\mathrm{TT}}$ is set to $\sigma$, and both $b_{\mathrm{HH}}$ and $b_{\mathrm{HT}}$ are set to $0.95\sigma$ (1, 2, 4), where $\sigma$ and $0.95\sigma$ are the diameter of a tail and a head bead, respectively.

**Bonded interactions**
FENE bonds potential presented between each successive pair of beads is given by

$$V_{\mathrm{bond}}(r) = -\frac{1}{2}k_{\mathrm{bond}}r_{\infty}^{2}\log\left[1-\left(\frac{r}{r_{\infty}}\right)^{2}\right],$$

where the bond stiffness $k_{\mathrm{bond}} = 30\lambda/\sigma^2$ and the divergence length $r_\infty$ is $1.5\sigma$.

A Hookean spring ensures that the preferred distance between the centers of the first and the last bead is $6\sigma$, to restrict bending fluctuations. The potential is

$$V_{\mathrm{bend}}(r) = \frac{1}{2}k_{\mathrm{bend}}(r-6\sigma)^2,$$

where $k_{bend} = 10\lambda/\sigma^2$.

**Simulation procedures**

For simulations using externally applied forces to press the vesicle membranes together, we placed 2 equilibrated vesicles (1 μs equilibration) within a cylinder extending along the z-axis with a radius $R_{\text{cyl}} = 30$ nm (Fig. 1B). The cylindrical wall interacts with lipid beads via a repulsive potential,

$$V_{\text{cyl}}(r) = \begin{cases} 4\epsilon_{\text{wall}} \left(\frac{\sigma_{\text{wall}}}{r}\right)^{12}, & r < r_{\text{cut}} \\ 0, & r \geq r_{\text{cut}} \end{cases}$$

where $\epsilon_{\text{wall}} = 0.06 k_B T$; $\sigma_{\text{wall}} = 0.44$ nm, equal to the lipid tail bead radius, and $r_{\text{cut}} = 2b_{\text{tt}} \simeq 1.8$ nm. The second vesicle is a copy of the first vesicle after a 3 μs equilibration period and was placed, so the outer leaflets of vesicle membranes were 4 nm apart initially. Then we applied a constant force of magnitude $f_{\text{const}}$ to every lipid bead to bring the two vesicles together.

For all group of SNARE-mediated simulations and TLT-mediated simulations, we build up two 50 nm vesicles made by crystal lipids and each SNARE-like fusogens were evenly placed in a 10 nm ring whose center is on the line between two vesicle centers. We equilibrated the whole system by 1 $\mu$s with the time step 1/100$^{\text{th}}$ of that in actual simulations. Then we ran 1 ms for TLT-mediated fusion simulations and ran 1.36 ms for SNARE-mediated fusion simulations.

**SNARE-ring radius measurements**

We measured the radius of the SNARE-ring by extracting the position of the LD N-terminus points and then calculating the mean relative distance between every LD N-terminus point and the center of the region surrounding by LD N-terminus points on the XY-plane. This means relative distance is the effective radius of the SNARE-ring.

#### Supplementary figures

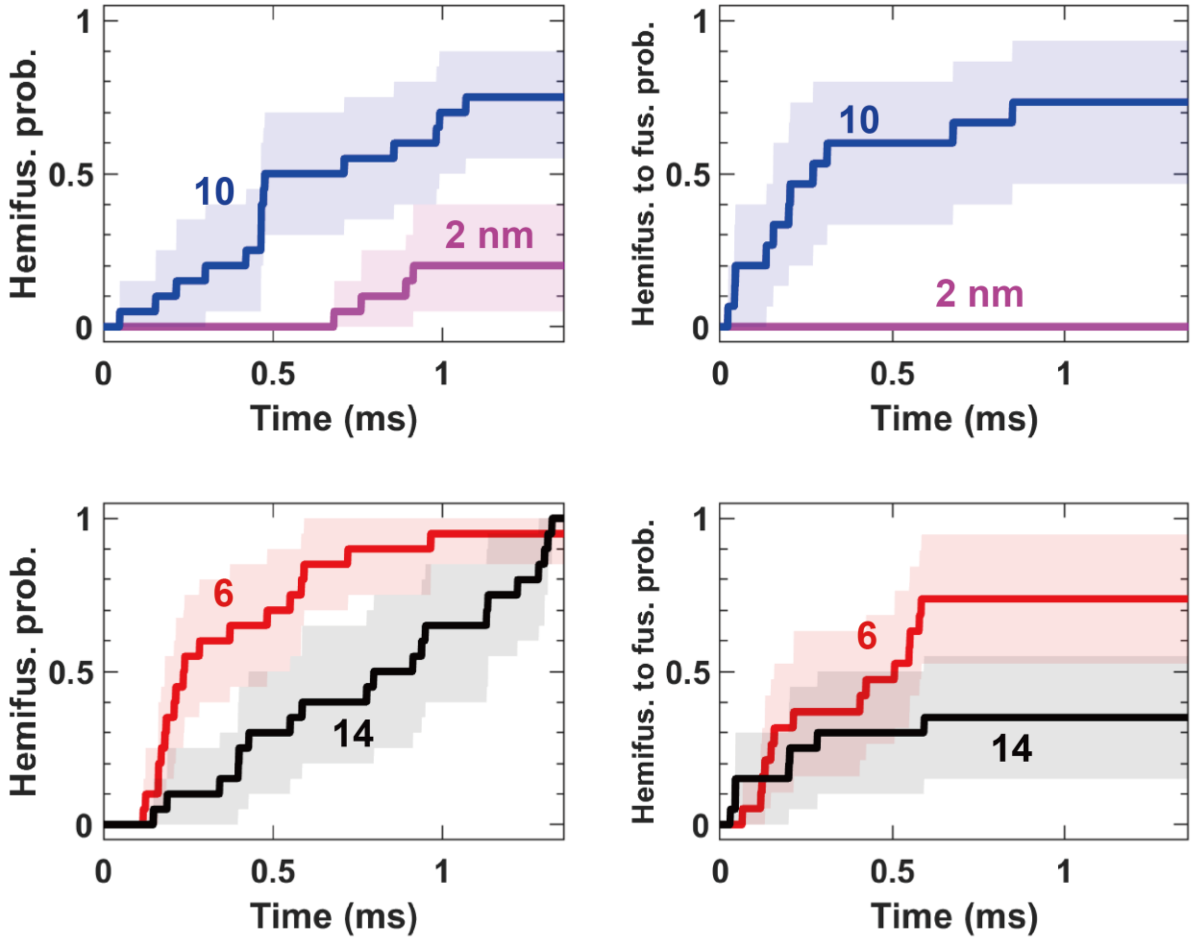


**Figure S1.** Cumulative of the fraction of simulations (n=20) achieved hemifusions (left) and fusion after hemifusion happened (right) using mutants with different SNARE motif lengths (6 SNAREs). The shaded regions were the 95% confidence interval computed by the boot-strapping method.

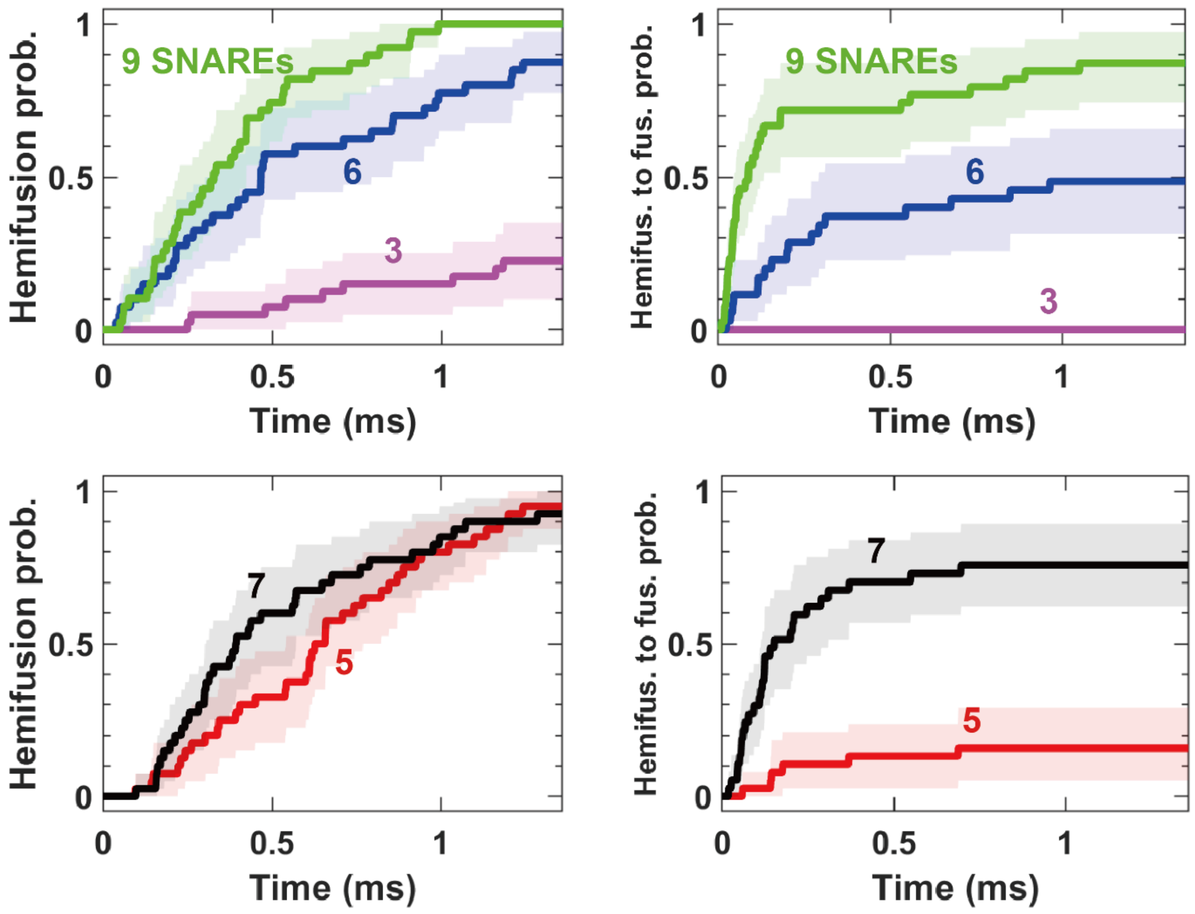


**Figure S2.** Cumulative of the fraction of simulations (n=40) achieved hemifusions (left) and fusion after hemifusion happened (right) using different numbers of SNAREs (SNARE length: 10 nm). The shaded regions were the 95% confidence interval computed by the boot-strapping method.

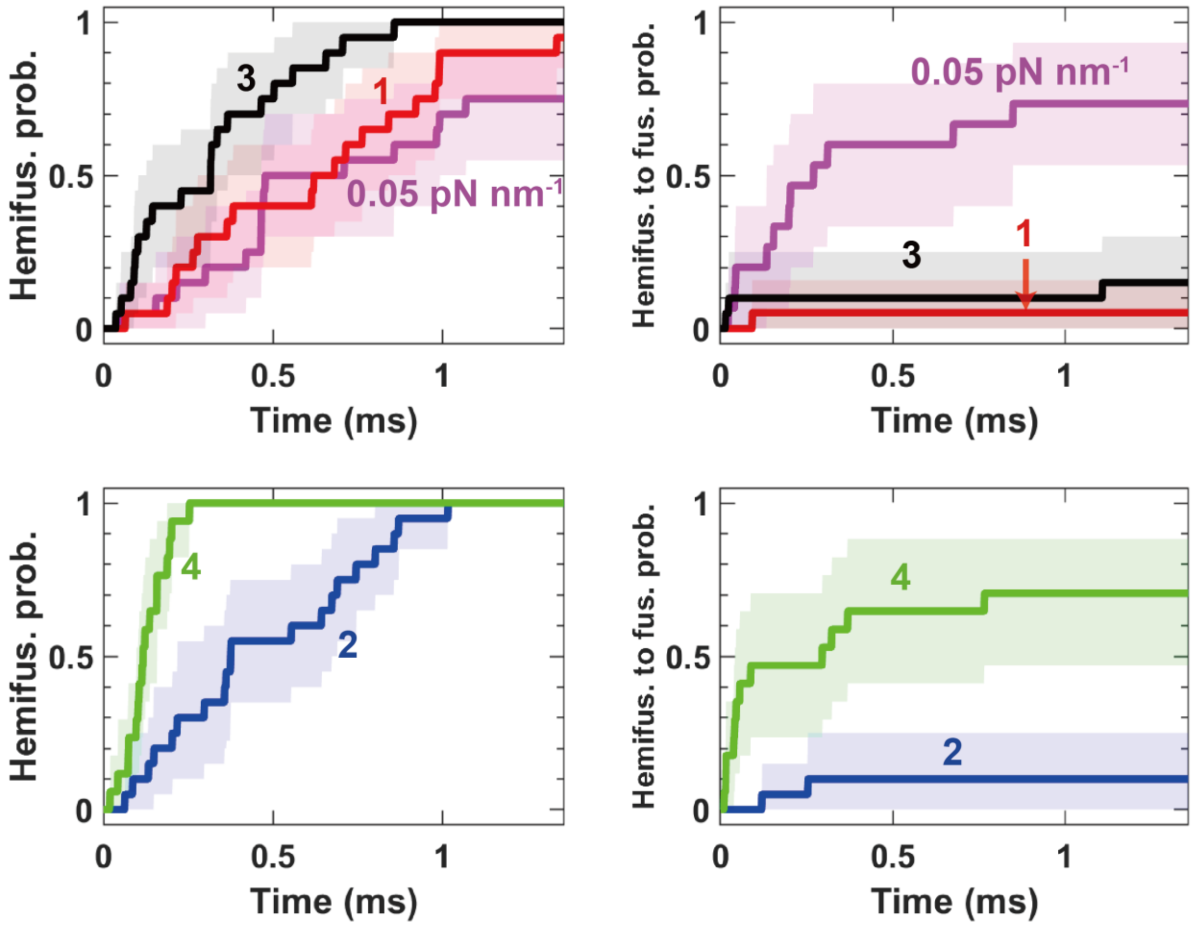


**Figure S3.** Cumulative of the fraction of simulations (n=20) achieved hemifusions (left) and fusion after hemifusion happened (right) under different vesicle tensions. The shaded regions were the 95% confidence interval computed by the boot-strapping method.

| Symbol | Meaning | Value | Legend |
|---|---|---|---|
| $\sigma$ | Length scale in the Cooke model | 0.88 nm | (A) |
| $\epsilon$ | Unit of energy in the Cooke model | 0.6 $k_B$T | (B) |
| $b_{hh}, b_{ht}$ | Head-head, head-tail repulsive length scale | $0.95\sigma$ | (B) |
| $b_{tt}$ | Tail-tail repulsive length scale | $\sigma$ | (B) |
| $w_c$ | Width of attractive potential between tail beads $V_{attr}$ | 1.4 nm | (C) |
| $D_{SNARE}$ | Diameter of the SNARE bead | 2 nm | (D) |
| $D_{TMD}$ | Diameter of the SNARE bead | 1 nm | (D) |
| $L_{TMD}$ | Length of the SNARE transmembrane domains | 3 nm | (D) |
| $L_{SNARE}$ | Length of the SNARE motif | 2, 6, 10, 14 nm | (D, E) |
| $T_{zip}$ | SNARE zippering force | 6-30 pN | (F) |
| $w_{TMD,TMD}$ | Width of attractive potential between TMD beads $V_{attr}$ | $1.6D_{TMD}$ | (G) |
| $\epsilon_{TMD,t}$ | Unit of energy in attractive potential between tail beads and TMD hydrophobic beads | $3\epsilon$ | (H) |
| $\epsilon_{TMD,h}$ | Unit of energy in attractive potential between head beads and either TMD hydrophilic beads or C-terminus LD beads | $2\epsilon$ | (H) |
| $L_x, L_y, L_z$ | Side lengths of the simulated bilayer | $\sim 88, 88, 123$ nm | |
| $\Delta t$ | Simulation timestep | 0.068 ns, 0.68 ps | (I) |

**Table S1.** Parameters of the coarse-grained force field used to represent lipid molecules developed in (1-3) and other simulation parameters for SNARE-like fusogens.

(A) Obtained by setting the measured bilayer thickness in the simulation to a typical experimentally measured value of 5 nm (5).
(B) Obtained from ref. (2).
(C) Set equal to 1.6 σ as in (1-3).
(D) Obtained from ref. (6).
(E) $L_{SNARE} = 10$ nm is the wide type SNARE, and the other lengths are mutants.
(F) $T_{zip} = 18$ pN is obtained from ref. (7) and the other zippering forces are mutants.
(G) Set equal to 1.6 $D_{TMD}$ similar to (1-3).
(H) The attraction potential parameters between TMD beads and lipid beads were set to prevent TMD pullout.
(I) A timestep of 0.068 ns was used for all of our measurements, except at equilibration, the timestep was 0.68 ps.

### Supplementary References


1. I. R. Cooke, K. Kremer, M. Deserno, Tunable generic model for fluid bilayer membranes. *Phys Rev E Stat Nonlin Soft Matter Phys* **72**, 011506 (2005).
2. G. Illya, M. Deserno, Coarse-grained simulation studies of peptide-induced pore formation. *Biophys J* **95**, 4163-4173 (2008).
3. I. R. Cooke, M. Deserno, Solvent-free model for self-assembling fluid bilayer membranes: stabilization of the fluid phase based on broad attractive tail potentials. *J Chem Phys* **123**, 224710 (2005).
4. I. R. Cooke, M. Deserno, Solvent-free model for self-assembling fluid bilayer membranes: Stabilization of the fluid phase based on broad attractive tail potentials. *J Chem Phys* **123** (2005).
5. B. A. Lewis, D. M. Engelman, Lipid bilayer thickness varies linearly with acyl chain length in fluid phosphatidylcholine vesicles. *J Mol Biol* **166**, 211-217 (1983).
6. A. Stein, G. Weber, M. C. Wahl, R. Jahn, Helical extension of the neuronal SNARE complex into the membrane. *Nature* **460**, 525-U105 (2009).
7. Y. Gao *et al.*, Single Reconstituted Neuronal SNARE Complexes Zipper in Three Distinct Stages. *Science* **337**, 1340-1343 (2012).
